# Solid-Phase Extraction Capture (SPEC) in nanoliter volumes for fast, robust and ultra-sensitive proteomics

**DOI:** 10.1101/2025.05.31.657165

**Authors:** Tim Heymann, Denys Oliinyk, Lukas Henneberg, Edwin Rodriguez, Anastasiya Bardziukova, Marc Oeller, Marta Murgia, Jesper V. Olsen, Stoyan Stoychev, Nicolai Bache, Matthias Mann

**Affiliations:** Department of Proteomics and Signal Transduction, Max Planck Institute of Biochemistry, Martinsried, Germany; Department of Biomedical Sciences, University of Padova, 35131, Padua, Italy; Novo Nordisk Foundation Center for Protein Research, University of Copenhagen, Copenhagen, Denmark; Evosep Biosystems, Odense, Denmark

## Abstract

Sample preparation remains a critical bottleneck in mass spectrometry (MS)-based proteomics, particularly for limited sample amounts where surface adsorption and dilution cause substantial losses. Here, we present Solid-Phase Extraction Capture (SPEC), a workflow that confines protein processing to nanoliter volumes within ion-exchange or C18 matrix inside a pipette tip. This achieves near-complete proteolysis within 5 minutes instead of hours and maintains full compatibility with strong detergents without cleanup steps, enabling effective lysis of challenging samples. From 200 ng FFPE tissue, SPEC achieves proteome depth and reproducibility exceeding conventional bulk protocols using 100 µg, critical when sample is irreplaceable. The modular two-tip configuration enables on-tip chemical modifications for mTRAQ labeling and fractionation, while integration with enrichment workflows yields 2-fold improved glycopeptide identifications from plasma and 3-fold enhanced ubiquitin remnant identification at low amounts. SPEC enables nanoPhos for cell-type resolved tissue phosphoproteomics and provides a universal platform for proteomics sample preparation.

## Introduction

Sample preparation has emerged as a principal limitation in MS-based proteomics, particularly as instrumentation advances have improved sensitivity to allow measurement of single cells and spatially defined tissue regions^1,2^. While modern mass spectrometers can detect and quantify peptides at attomole amounts, the journey from biological sample to injection-ready peptides remains fraught with losses that compound at each step: lysis, reduction, alkylation, digestion, and cleanup. For bulk samples measured in micrograms, these losses are tolerable, however, for emerging applications that define the field’s trajectory, they become prohibitive.

The proteomics community has responded with increasingly sophisticated sample preparation strategies. Filter-aided sample preparation (FASP) enabled SDS-based lysis by retaining proteins on molecular weight cutoff membranes during buffer exchange^3^. The iST method consolidated processing steps into an enclosed filter-based format^4^. SP3 and the related PAC protocol captured proteins on functionalized magnetic beads, reducing transfers and enabling automation^5–7^. Tip-based approaches, including STAGE tips and one-pot reactors, combined solid-phase extraction with enzymatic processing in miniaturized formats^8–10^. Each advance has improved some aspect of the workflow - detergent compatibility, automation potential, or reduced handling - yet all share the fundamental constraint that they operate at microliter volumes where the physics of surface adsorption and enzyme kinetics work against efficient processing of limited samples^11,12^.

We reasoned that confining both sample and enzyme to nanoliter volumes within a solid-phase matrix would fundamentally alter these dynamics^12,13^. By binding proteins to an ion-exchange or reversed-phase resin and introducing enzyme into this confined space, we could achieve local concentrations orders of magnitude higher than solution-phase methods while minimizing surface exposure. Here we present Solid-Phase Extraction Capture (SPEC), a workflow built on this principle. SPEC processes proteins within minute volumes inside pipette tips packed with solid phases, achieving near-complete digestion in minutes rather than hours while maintaining full compatibility with strong detergents. The resulting peptides transfer directly to Evotips for liquid chromatography mass spectrometry (LC-MS) analysis without intermediate steps.

We demonstrate that SPEC provides a universal platform for proteomics sample preparation: it handles fresh lysates and formalin-fixed paraffin-embedded (FFPE) tissue with equal facility – from ug amounts to single cell equivalents. Its modular design enables on-tip chemical modifications including isotope labeling and pH-based fractionation, while integration with enrichment workflows substantially improves glycopeptide and ubiquitin remnant identification. Importantly, SPEC serves as the foundation technology for nanoPhos^14^, where its efficiency enables ultrasensitive and cell type-resolved phosphoproteomics from spatially defined tissue regions. We anticipate this workflow will become a standard for proteomics sample preparation across the full spectrum of applications, from high-throughput clinical studies to spatial single-cell analyses.

## Results

### Nanoliter confinement enables rapid and complete proteolysis

The defining principle of SPEC is physical confinement of proteins to nanoliter-scale volumes for all processing steps (**Fig. 1a**). We load reduced and alkylated protein lysates onto common solid-phase tips - either C18 for reversed-phase capture or ion-exchange resins (SCX or SAX) - where they concentrate within the resin bed. Proteolytic enzymes introduced into this confined space achieve effective concentrations far exceeding those possible in conventional solution-phase digestion.

**Figure 1.**
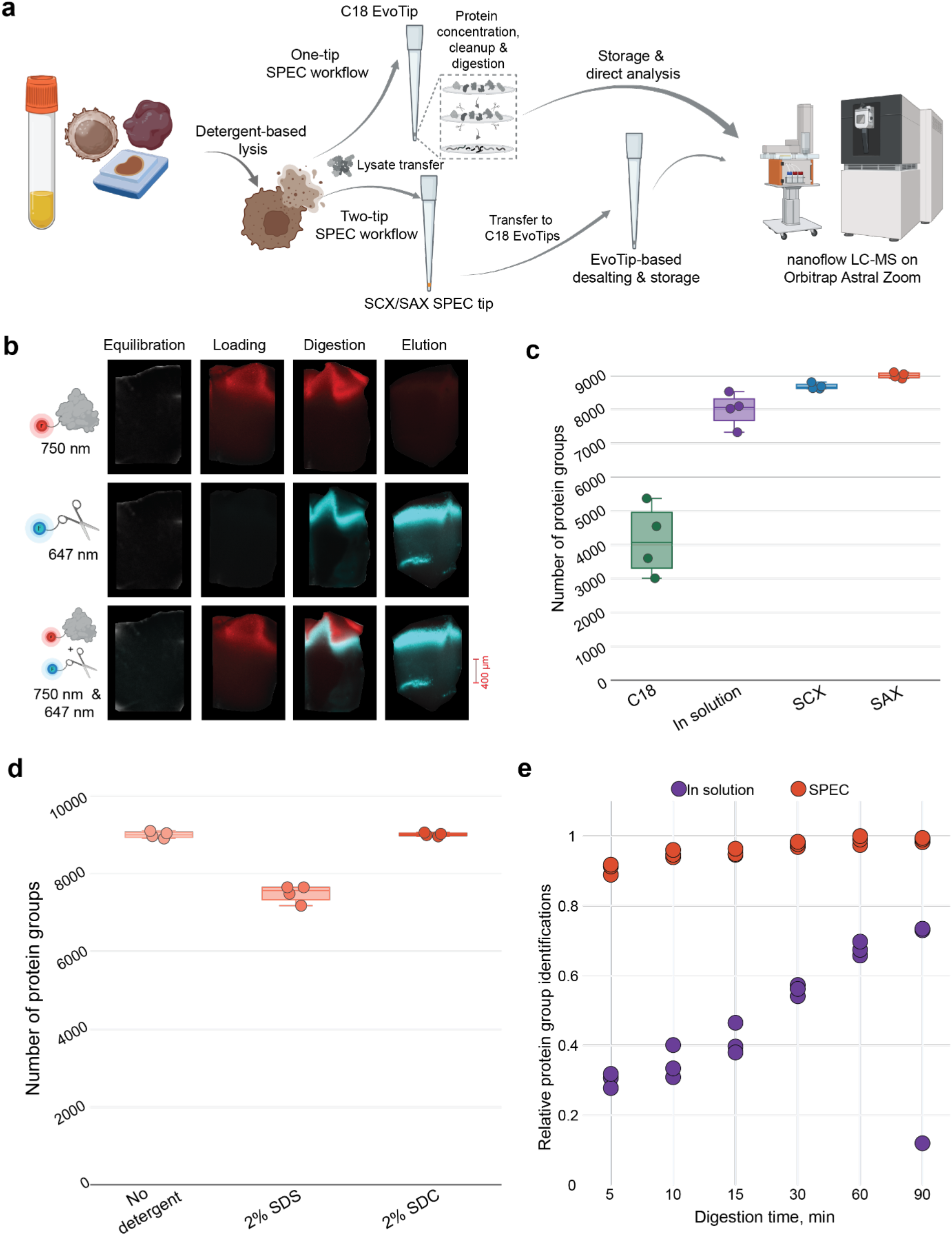
SPEC confines protein processing to nanoliter volumes for efficient proteolysis. **a** Overview of the SPEC workflow. Protein samples from diverse sources (body fluids, cells, and FFPE tissue) undergo detergent-based lysis, followed by processing through either a one-tip configuration using C18 EvoTips directly or a two-tip configuration where proteins are loaded onto SCX/SAX SPEC tips for digestion, then transferred to C18 EvoTips for desalting and storage until nanoflow LC-MS analysis on an Orbitrap Astral Zoom. **b** Fluorescence imaging of SAX-SPEC tips throughout the workflow. K562 lysate labeled with NHS-Alexa750 (red, 750 nm; top row) and trypsin/LysC labeled with NHS-Alexa647 (cyan, 647 nm; middle row). Bottom row shows the merged channels for lysate and protease. Scale bar, 400 μm. White arrows indicate the penetration depth of approximately 200 µm of digested protein and enzyme at the end of digestion in the material. **c** Protein group identifications from 200 ng K562 lysate comparing four conditions: C18 single-tip, in-solution digestion (on-top-of-tip), SCX-SPEC and SAX-SPEC. Box plots show replicate distributions with individual data points (n = 4 per condition). **d** Protein group identifications from 200 ng K562 lysate processed with SAX-SPEC using no detergent, 2% SDS, or 2% SDC. Box plots show replicate distributions with individual data points (n = 4 per condition). **e** Protein group identifications (normalized to maximum) for SAX-SPEC (red) versus in-solution digestion (with SDB-RPS cleanup) (purple) from 5 to 90 minutes. Box plots show replicate distributions with individual data points (n = 3 per condition).

While single-tip SPEC using commercial Evotips (C18 material) is convenient, we developed a two-tip configuration to optimize each processing step independently (**Fig. 1a, c**). In this format, proteins bind and digest on a SPEC tip containing SAX or SCX resin, then peptides elute directly onto a standard C18 Evotip for desalting and storage until LC-MS analysis. This separation allows us to use the optimal chemistry for digestion (SAX) at pH 8.5 while maintaining compatibility with the Evosep One LC system.

We implemented this configuration using 3D-printed adapters that position SPEC tips directly above Evotips in 96-well formats (**Suppl. Fig. 1a**). Centrifugation transfers peptides from the digestion tip to the desalting tip without manual intervention, enabling parallel processing of entire plates. This design integrates seamlessly with liquid handling robots for fully automated workflows.

To visualize this confinement directly, we fluorescently labeled K562 protein lysate and separately trypsin/LysC, then imaged SAX-SPEC tips throughout the workflow using fluorescence microscopy (**Fig. 1b**). After equilibration, the SAX material showed no fluorescence. Upon loading, 200 ng proteins concentrated sharply at the tip of the resin bed, forming a well-defined band. As proteins are confined to the top 200 µm of the solid phase at the end of the digestion as indicated by the white arrows, we estimate the digestion volume to be well below 200 nL, depending on the sample amount. When enzyme was introduced, it colocalized with the protein band, confirming that digestion occurs within this confined nanoliter volume rather than in the bulk liquid above the resin. After elution, the lysate signal disappears while most of the protease signal is retained on the resin, indicating complete digestion and peptide release without co-elution of enzymes.

We compared SPEC across three solid-phase chemistries - C18, SCX, and SAX - against in-solution digestion of 200 ng K562 lysate with readily available and stable buffers at a matching pH without further optimization (**Fig. 1c**). SAX-based SPEC performed best achieving 9,100 protein groups with the Evosep Whisper Zoom 80 samples per day (SPD) LC gradient and narrow-window data-independent (nDIA) on an Orbitrap Astral^15^, compared to 4,100 for C18, 8,500 for in-solution and 8,800 for SCX based digestion. Given SAX-SPEC’s superior performance and sample recovery (**Suppl. Fig. 1b, c**), we evaluated its compatibility with common lysis detergents. SAX-SPEC maintained optimal performance with 2% SDC (9,700 protein groups), comparable to no detergent in the input, while 2% SDS reduced identifications to 7,600 protein groups (**Fig. 1d**). This compatibility with strong detergents, preferably SDC, eliminates cleanup steps that introduce losses in other workflows.

To study digestion kinetics, we compared SAX-based SPEC to conventional in-solution digestion with SDB-RPS cleanup across time points from 5 to 90 minutes using 200 ng HeLa lysate (**Fig. 1e**). SPEC reached approximately 95% of maximum protein group identifications within 5 minutes, with essentially complete digestion by 30 minutes. By contrast, in-solution digestion achieved only 30% completion at 5 minutes and required > 90 minutes to approach SPEC’s 5-minute performance. This 20-fold acceleration is not merely convenient for workflow timing - it demonstrates that the confined geometry creates fundamentally more efficient conditions for proteolysis. Nevertheless, in order to maintain robust digestion efficiency for harder to digest samples, we used a digestion time of 1h for all subsequent experiments.

### SPEC maximizes sensitivity and enables on-tip modifications

To quantify SPEC’s performance at low sample inputs, we processed K562 lysate across a dilution series from 1 µg to 0.25 ng - the approximate protein content of a single mammalian cell (**Fig. 2a**). We compared SPEC to in-solution digest on top of the SPEC tip and an alternative concentration method – Protein Aggregation Capture (PAC), where proteins are precipitated on the surface of hydroxyl beads^7^. SPEC outperformed PAC and in-solution digestion across the entire range, with the advantage becoming most pronounced at the lowest amounts. At single-cell-equivalent input (0.25 ng), SPEC identified approximately 13% more protein groups as PAC and about 2-fold compared to in-solution. In both cases, a substantially higher fraction was quantified at CV <20% (**Suppl. Fig. 2a**). This advantage was maintained across the entire input range that we tested, but most pronounced at 50 ng and below. Our results demonstrate the consistently higher sample recovery of SPEC compared to in-solution as well as PAC at an equal digestion time of 1h. This is also reflected in the median precursor sum that is consistently higher for SPEC until background peptides from the protease start becoming a major contributor - at around 1 ng lysate input (**Suppl. Fig 2b**).

**Figure 2.**
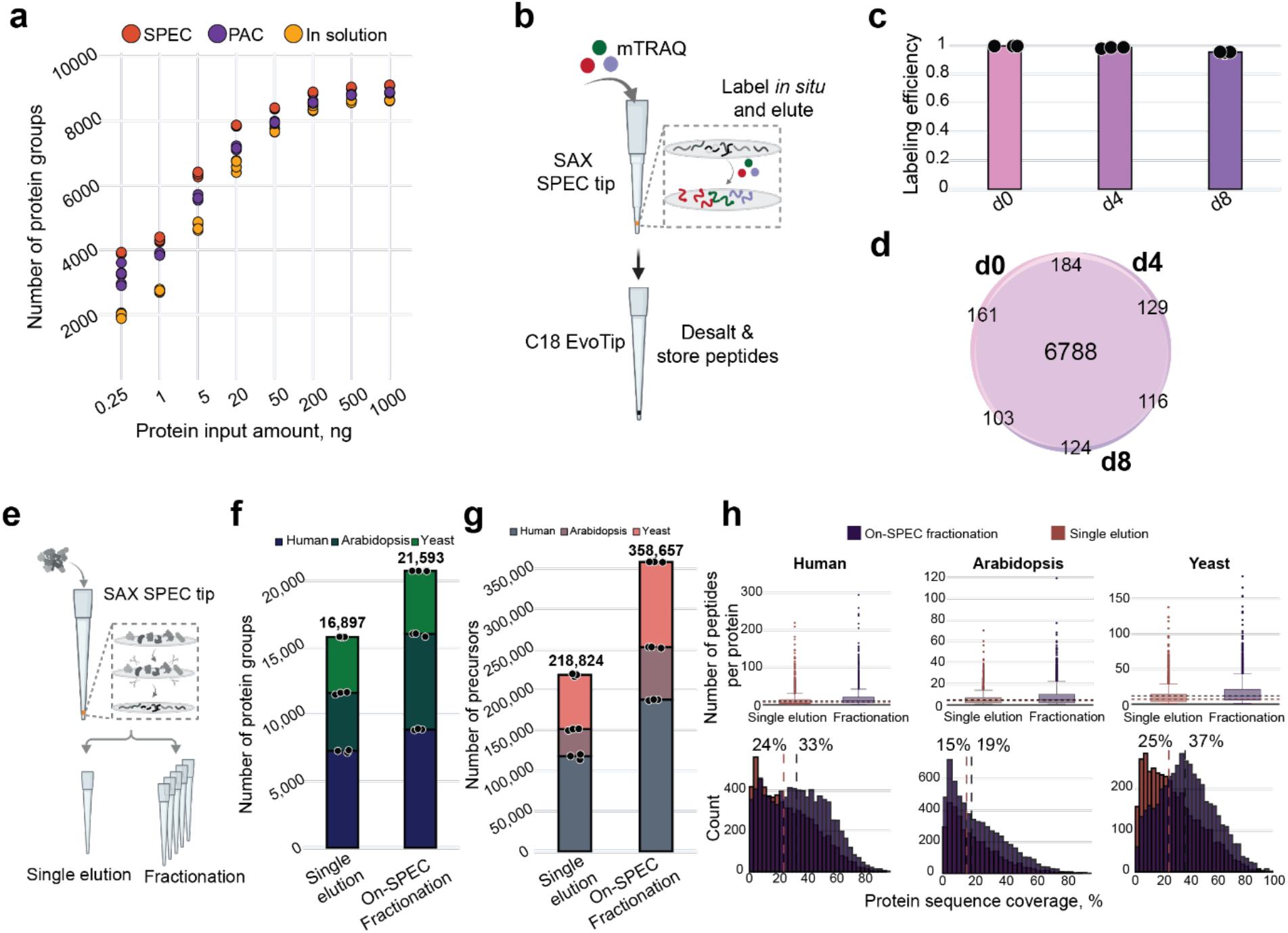
SPEC achieves high sample recoveries and allows on-tip manipulation of peptides and proteins to increase proteomic depths and throughput. **a** Protein group identifications across an input titration series from 0.25 ng (single-cell equivalent) to 1,000 ng K562 lysate comparing SPEC (red), PAC (purple) and in-solution (yellow) (on-top-of-tip) (n = 3 per condition). **b** Schematic of on-tip mTRAQ labeling workflow. After digestion on SAX SPEC tips, mTRAQ reagent (three channels: d0, d4, d8) is added directly to the tip for in situ labeling. Labeled peptides elute onto C18 EvoTips for desalting and storage. **c** Labeling efficiency for the three mTRAQ channels (d0, d4, d8) on SAX SPEC tips using 200 ng K562 lysate with median (bars) and individual replicates (n = 3 per condition) shown. **d** Venn diagram showing overlap between d0, d4 and d8 mTRAQ channels. **e** Schematic of on-SPEC fractionation workflow. Peptides bound to SAX SPEC tips after digestion can be eluted either in a single step or through salt gradient-based fractionation. **f** Protein group identifications from three-proteome mixture (human, Arabidopsis, yeast) comparing single elution versus on-SPEC fractionation. Stacked bars show species-specific contributions with median (bars) and individual replicates (n = 3 per condition). **g** Same as **f** but showing precursor identifications. **h** Histograms comparing peptide coverage metrics between on-SPEC fractionation (purple) and single elution (red) for each organism (Yeast, Human, Arabidopsis). Top row: distribution of peptides per protein. Bottom row: protein sequence coverage distributions with median values indicated. Based on results shown for **f** and **g**.

The confinement of proteins to nanoliter-scale volumes has benefits beyond digestion. SAX-SPEC enables on-tip chemical modifications. After digestion but before elution, the peptides remain bound to the SAX resin in a defined, accessible volume. We exploited this for mTRAQ isotope labeling (**Fig. 2b**) by replacing the nucleophile CAPS loading buffer with an unreactive sodium carbonate buffer at the same pH. Adding reagent to the SPEC tip after digestion achieved >95% labeling efficiency for 200 ng of input lysate across all three channels (d0, d4, d8) with 7,000 protein groups identified per channel and >97% overlap between channels (**Fig. 2c,d**). Note that this is at least on par with our recent labeling results for single cell proteomics in which we used DDA to assess the degree of completeness^16^ (**Methods**). This on-tip labeling eliminates the peptide cleanup steps typically required between digestion and labeling, further reducing sample losses. This capability opens the door to multiplexed quantification directly from limited samples, enabling comparative studies that would otherwise require pooling or compromise on replicates.

SAX is a material classically used for fractionation of complex biological samples to reduce complexity and improve proteome coverage. SPEC utilizing SAX enables the combination of digestion, cleanup and fractionation in one step (**Fig. 2e**). Similar to earlier StageTip-based fractionation approaches^17,18^ we applied a salt gradient directly to the SAX SPEC tip rather than eluting all peptides in a single step. We collected a three-proteome mixture (human, Arabidopsis, yeast) in five sequential fractions onto separate Evotips. On-SPEC fractionation increased protein group identifications from 16,900 to 21,600 (28% improvement, **Fig. 2f**) and peptide precursor identifications from 218,800 to 358,600 (64% improvement, **Fig. 2g**) starting with just 2.5 ug of input lysate. This gain was accompanied by improved peptide counts per protein and protein sequence coverage across all three organisms (**Fig. 2h**), demonstrating that the fractionation accesses peptides missed in single-shot analysis without requiring additional sample input. Importantly, this integrated digestion-fractionation workflow requires no additional sample handling compared to standard SPEC, making deep proteome coverage accessible even when material is limiting.

### SPEC achieves deep proteomes from limited tissue samples

A stringent test of any sample preparation method is performance on formalin-fixed paraffin-embedded (FFPE) tissue, where crosslinking, embedding media, and limited material availability compound the usual challenges. We processed 200 ng of mouse liver FFPE lysate with SPEC and compared it to both PAC and in-solution digestion (**Fig. 3a**).

**Figure 3.**
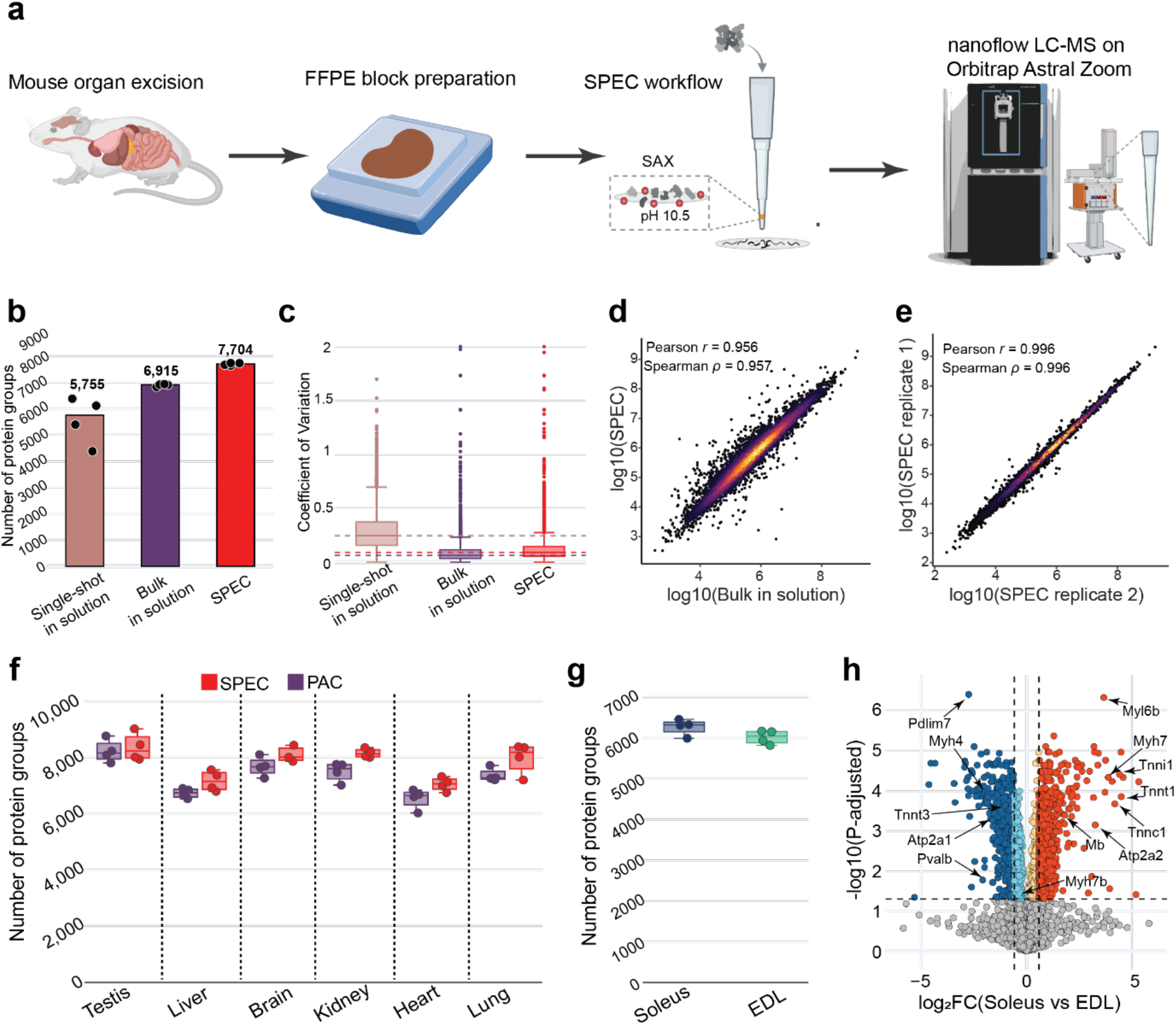
SPEC enables deep proteomes from FFPE tissue and distinguishes muscle fiber types. **a** Workflow for processing mouse FFPE tissue. After mouse organ excision, tissues undergo FFPE block preparation. Deparaffinized samples are lysed and processed through SAX-SPEC at pH 10.5, followed by nanoflow LC-MS analysis on an Orbitrap Astral Zoom. **b** Protein group identifications from mouse liver FFPE comparing three conditions: single-shot 200 ng in-solution, bulk 100 μg in-solution, and 200 ng SAX-SPEC (n = 4 per condition). **c** Coefficient of variation (CV) comparison for the results shown in **b.** Dashed lines indicates median CV for each condition. **d** Correlation of protein intensities between SPEC and bulk in-solution digestion with Pearson *r* and Spearman *ρ* indicated. Based on the data shown in **b. e** Technical replicate correlation for SPEC with Pearson r and Spearman ρ indicated. Based on the data shown in **b. f** Protein group identifications across six mouse FFPE tissues (testis, liver, brain, kidney, heart, lung) comparing PAC (purple) and SPEC (red). Box plots show replicate distributions with individual data points (n = 4 per condition). **g** Protein group identifications from freeze-dried mouse skeletal muscle fiber bundles processed with SAX-SPEC. Box plots show replicate distributions with individual data points (n = 4 per condition). **h** Volcano plot (P. Adj. < 0.05; |log2FC| > 0.585) of soleus versus EDL proteomes showing significantly enriched proteins. Dashed lines indicate significance thresholds.

From 200 ng input, SPEC achieved 7,700 protein groups per replicate and coefficient of variation comparable to a 100 ug bulk in-solution digest (median CV = 0.11, **Fig. 3b, c**). Single-shot in-solution digestion of the same 200 ng yielded only 5,750 protein groups per replicate, while even bulk in-solution using 100 µg achieved only 6,500 protein groups per replicate. This demonstrates that for samples where material is fundamentally limited, SPEC recovers proteome depth that conventional methods cannot access regardless of input amount. This difference is even more evident comparing the median precursor intensity, where SPEC achieves an 8-fold gain over in-solution digest at the same input **(Suppl. Fig. 3a)**. The Pearson correlation between SPEC and bulk in-solution reached 0.96, confirming that SPEC does not introduce systematic biases (**Fig. 3d**). Technical replicates of SPEC showed near-perfect correlation (r = 0.996, **Fig. 3e**).

We extended this analysis across six FFPE mouse tissues: testis, liver, brain, kidney, heart, and lung (**Fig. 3f**). SPEC consistently achieved 7,000–10,000 protein groups from 200 ng input across all tissues, substantially outperforming PAC digestion on matched samples. This is due to SPEC’s significantly higher sample recovery across all tissue types with up to two times higher median precursor sum intensities at the same sample input (**Suppl. Fig. 3b**). Principal component analysis (PCA) separated tissues cleanly by organ identity (**Suppl. Fig. 3c**). This biological separation confirms that SPEC preserves the quantitative proteome differences that define tissue identity.

Next, we focused on a tissue presenting exceptional challenges for proteomics: Skeletal muscle proteomes are dominated by highly abundant contractile proteins while functionally important regulatory proteins span orders of magnitude lower abundance, and the dense extracellular matrix resists digestion. Our SPEC-based analysis of soleus and EDL fiber bundles achieved >6,000 protein groups from each (**Fig. 3g**). Differential expression analysis showed soleus, a slow-twitch oxidative muscle, enriched for MYH7 (slow myosin) and ATP2A2 (SERCA2), while EDL, a fast-twitch glycolytic muscle, was enriched for MYH4 (fast myosin IIb) and ATP2A1 (SERCA1). These represent among the deepest untargeted proteomes reported for mouse skeletal muscle, achieved from 200 ng of freeze-dried fiber material. Differentially regulated proteins are spread across the entire dynamic range of the detected muscle proteome demonstrating biologically meaningful data gained by improved sample processing (**Suppl. Fig. 3d**).

The clear molecular distinction between fiber types, including metabolic enzymes, calcium handling proteins, and sarcomeric components, demonstrates that SPEC preserves the biological signatures essential for understanding muscle physiology and disease.

### SPEC integrates with downstream enrichment workflows

We reasoned that beyond direct proteome analysis, enrichment for post-translational modifications may benefit significantly from SPEC’s efficient digestion, high sample recovery and clean peptide output. First, we tested this for plasma glycoproteomics, a technically demanding application where the combination of high dynamic range, glycan heterogeneity and limited sample availability has impeded progress (**Fig. 4a**).

**Figure 4.**
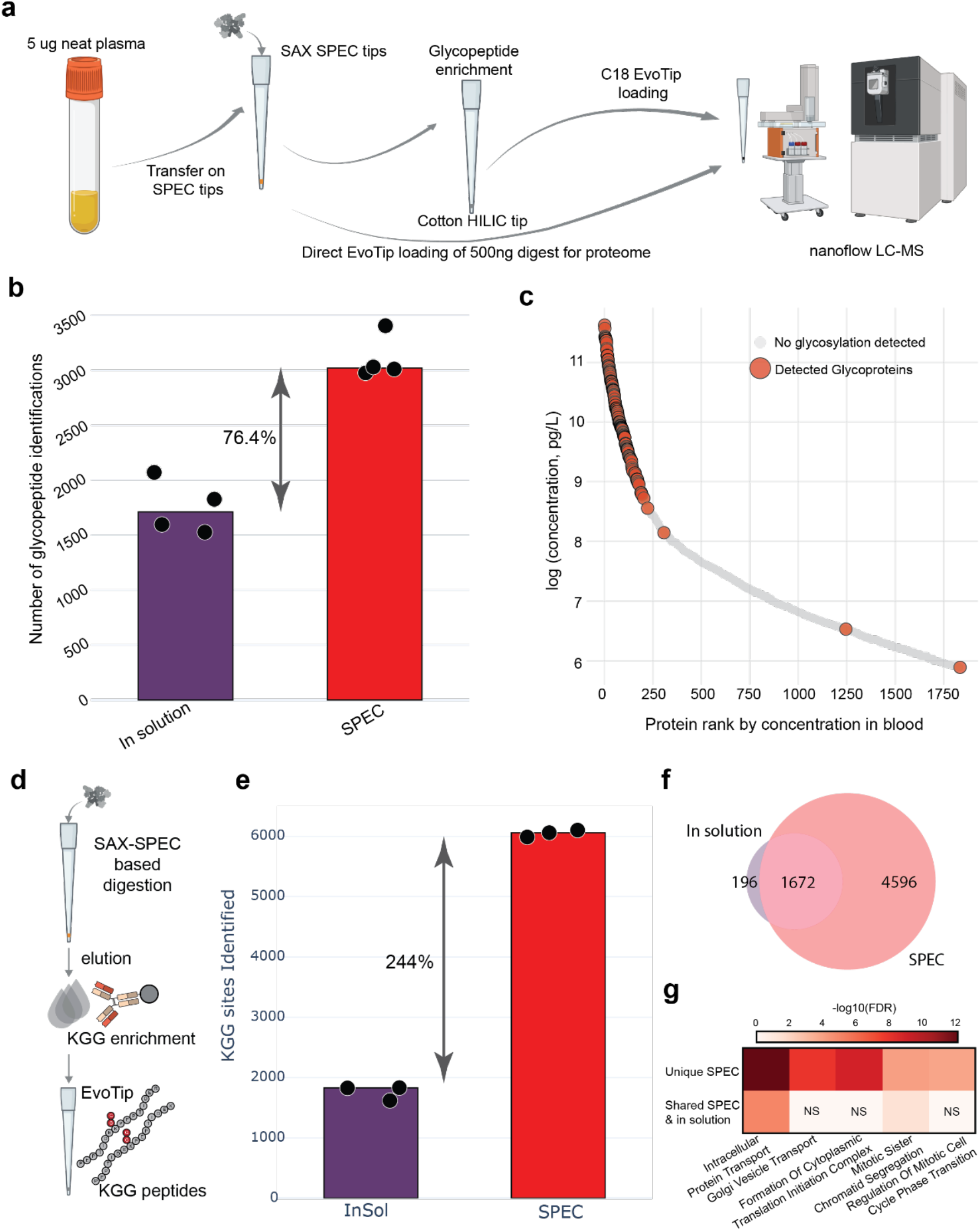
SPEC integrates with downstream enrichment for glycoproteomics and ubiquitin remnant profiling. **a** Workflow for plasma glycoproteomics. Starting with 5 μg neat plasma, proteins are transferred onto SAX SPEC tips for concentration, cleanup, and digestion. A portion (500 ng) is loaded directly onto EvoTips for proteome analysis. The remainder undergoes glycopeptide enrichment via cotton HILIC tip, followed by C18 EvoTip loading and nanoflow LC-MS analysis. **b** Glycopeptide identifications from 5 μg plasma processed by in-solution or SPEC workflow prior to cotton HILIC tip enrichment, showing median (bars) and individual replicates (n = 4 per condition). **c** Protein rank plot of detected glycoproteins from **b** mapped to Human Blood Atlas plasma concentrations. **d** Workflow for SPEC-integrated ubiquitin remnant (KGG) profiling. After digestion on SAX SPEC tips, peptides are eluted and subjected to KGG enrichment using antibody-conjugated beads, yielding enriched diglycine-modified (ubiquitinated) peptides. **e** KGG peptide identifications from 5 μg cell lysate processed by in-solution or SPEC workflow prior to antibody-bead based KGG enrichment, showing median (bars) and individual replicates (n = 3 per condition). **f** Venn diagram showing overlap between in-solution and SPEC KGG identifications **g** GO term enrichment analysis for modified proteins, unique for SPEC and those that were identified in both SPEC-based and in solution-based conditions. We selected the GO terms shown in the panel to highlight the differences between the workflows (FDR < 0.05).

Starting with just 5 µg of neat plasma, we compared SPEC to in-solution digestion followed by cotton HILIC glycopeptide enrichment^19,20^. For the unfractionated proteome, SPEC and in-solution both identified about 1400 protein groups from 500 ng digest loaded in quadruplicate, with near-perfect Spearman correlation (r = 0.99) confirming that SPEC does not alter the plasma proteome profile (**Suppl. Fig 4a, b**).

The advantage emerged upon glycopeptide enrichment where SPEC-digested samples yielded ∼3,000 glycopeptide identifications compared to ∼1,700 for in-solution—a two-fold improvement from identical starting material (**Fig. 4b**). This gain reflects both the higher peptide purity from SPEC (facilitating enrichment) and the higher sample recovery (higher glycopeptide concentration). The identified glycoproteins spanned 5.5 orders of magnitude in plasma concentration according to the Human Blood Atlas (**Fig. 4c**), demonstrating that SPEC-based enrichment achieves comprehensive glycoproteome coverage from minimal clinical sample volumes. Interestingly, most of the detected 200 glycoproteins are concentrated in the upper end of the rank plot, presumably reflecting the fact that much of the functional and high abundance plasma proteome is glycosylated. This depth of coverage from less than a single µL of blood draw has immediate implications for glycoproteomics, where sample availability often constrains study design.

Next, we integrated SPEC into ubiquitin remnant (KGG) profiling^21–24^ (**Fig. 4d**), where antibody-based enrichment captures diglycine-modified peptides marking ubiquitination sites. From 5 µg of cell lysate, SPEC-based preparation yielded close to 6,000 KGG peptide identifications compared to 1,800 for in-solution digestion - a 3-fold improvement (**Fig. 4e**). Beyond 1,700 KGG shared peptides, SPEC identified 4,600 additional sites compared to only 200 unique to in-solution (**Fig. 4f**). This dramatic improvement demonstrates that SPEC’s efficient digestion and peptide recovery substantially expand access to the ubiquitinome, enabling deeper investigation of protein degradation and signaling pathways from limited samples.

While both workflows recovered core components of the ubiquitin– proteasome machinery, integration of SPEC with KGG enrichment revealed substantially broader biological pathway coverage. Gene Ontology analysis of ubiquitinated proteins uniquely identified using SPEC showed significant enrichment for processes extending beyond protein degradation, including Golgi vesicle transport, RNA processing, membrane organization, tRNA aminoacylation, and regulation of mitotic cell cycle phase transitions (**Fig. 4g**). These pathways represent distinct layers of cellular regulation encompassing protein synthesis, trafficking, and cell cycle control. In contrast, proteins shared between the two workflows did not exhibit comparable enrichment for these processes, likely reflecting the more limited analytical depth of in-solution digestion. Together, these results suggest that SPEC-based preparation not only increases ubiquitination site coverage but may also enable detection of additional, biologically meaningful pathways that remain inaccessible using conventional workflows.

## Discussion

SPEC addresses a fundamental physical problem in proteomics sample preparation. By confining proteins to nanoliter volumes within solid-phase tips, it transforms the kinetics of every processing step. Enzyme and substrate concentrations increase by orders of magnitude compared to conventional microliter-scale reactions. Surface-to-volume ratios decrease correspondingly, reducing adsorptive losses. The result is near-complete proteolysis in minutes rather than hours, with recovery efficiency that enables proteome depth from nanogram inputs previously requiring 100-fold more material.

Several features distinguish SPEC from prior tip-based methods. First, digestion occurs *within* the solid phase rather than in solution above it - proteins bound to the resin are directly accessible to enzymes and labeling reagents that diffuse into the confined matrix. Second, the two-tip configuration separates digestion chemistry and cleanup from peptide storage, allowing each to be independently optimized. Third, compatibility with strong detergents (SDS, SDC) eliminates the cleanup steps that introduce losses in other workflows. Fourth, the format integrates directly with the Evosep One LC system, enabling high-throughput operation without additional transfers.

The rapid digestion kinetics merit particular discussion. Completing proteolysis in an hour rather than overnight has obvious workflow advantages, but its greater significance is mechanistic: it demonstrates that the confined geometry creates conditions for efficient, complete digestion, even from very diluted samples. This has immediate practical implications for applications requiring rapid turnaround - clinical diagnostics, or experimental designs where prolonged incubation would alter the biological state of interest.

Beyond sample diversity, the two-tip configuration enables modular on-tip chemistry. After proteolysis, peptides remain bound to the SAX resin in a defined volume accessible for chemical modification. On-tip mTRAQ labeling achieved high efficiency across all three channels with over 97% peptide overlap, eliminating cleanup steps between digestion and labeling. The same principle enables extremely simple on-tip fractionation: stepwise salt elution from SAX tips increased protein identifications by 28% and precursor identifications by 64% compared to single-shot analysis, unlocking deeper proteome and isoform coverage when it is needed.

The consistency of SPEC performance across diverse sample types suggests the generality of the underlying physical principle. From six different mouse FFPE tissues, SPEC consistently achieved 7,000-10,000 protein groups from just 200 ng input-depths that bulk in-solution methods fail to reach even with 100 ug. Freeze-dried skeletal muscle fibers yielded over 6,000 proteins with clean separation of slow-twitch and fast-twitch markers, representing among the deepest untargeted muscle proteomes reported. Whatever the biological source, proteins captured on an optimized solid phase and digested in a confined volume behave predictably. This universality positions SPEC as a potential default method for proteomics sample preparation.

SPEC’s efficient digestion and clean peptide output substantially improve downstream enrichment workflows. For plasma glycoproteomics, SPEC doubled glycopeptide identifications compared to in-solution digestion from just 5 ug starting material, spanning 5.5 orders of magnitude in plasma protein concentration. Ubiquitin remnant profiling showed an even more dramatic three-fold improvement, with SPEC identifying nearly 5,000 KGG peptides versus 1,650 for conventional methods. These gains reflect probably both higher peptide purity facilitating enrichment and higher sample recovery increasing the concentration of the modified peptides. The enabling role of SPEC extends to phosphoproteomics: in our companion manuscript, nanoPhos combines SPEC-based digestion with zero-dead-volume phosphopeptide enrichment to identify over 4,000 phosphosites from 10 ng input, enabling spatially resolved phosphoproteomics from laser-microdissected tissue regions^14^.

As SPEC tips become commercially available, the barrier to adoption will be minimal. The workflow uses standard laboratory equipment—centrifuges, pipettes, and heating blocks—with no specialized instrumentation. Integration with liquid handling robots is straightforward, as demonstrated by our 96-well format processing. We expect SPEC to become a foundation for the next generation of proteomics applications, from clinical diagnostics requiring rapid results to spatial biology demanding maximum sensitivity from minimal tissue regions.

## Methods

### Cell culture and lysate preparation

Human epithelial carcinoma (HeLa, ATCC S3 subclone) and hTERT RPE-1 cells were cultured in DMEM containing 20 mM glutamine, 10% FBS, and 1% penicillin-streptomycin. Cells were tested routinely for mycoplasma contamination. For experiments, cells were cultured to 80% confluency, harvested with 0.25% trypsin/EDTA, washed twice with cold TBS, pelleted at 200×g for 10 min, snap-frozen, and stored until use. For lysis, pellets were resuspended in lysis buffer (100 mM Tris-HCl pH 8.5, 40 mM CAA, 10 mM TCEP, 0.01% DDM, 2% SDC) and boiled for 15 min at 95°C with mixing at 1,500 rpm. Lysates were subjected to high-energy tip sonication (10 pulses, 5 s on/5 s off, 20% duty cycle) and centrifuged at maximum speed for 5 min to remove debris. Protein concentration was determined by tryptophan assay, and samples were diluted to 1 µg/µL with 0.5% SDC in 100 mM Tris-HCl before storage at -80°C.

Commercial lysates were used for K562 (human), yeast (both from Promega), and Arabidopsis thaliana [Creative Biomart]. Lysates were diluted to 1 µg/µL in buffer containing 50 mM TEAB pH 8.5, 40 mM CAA, and 10 mM TCEP, incubated for 30 min at room temperature, aliquoted, and stored at -80°C.

### EvoTip Pure Sample loading

For loading of digested samples, Evotips Pure were prepared by washing with 20 µL EvoB (0.1% FA in ACN), priming in 1-propanol for 15 s, and washing with 20 µL EvoA (0.1% FA in water). The disk was wetted with 100 µL EvoA to keep it wet for the full SPEC elution process. Samples were then loaded by pipetting or by direct elution from SPEC tips. After loading, tips were washed with 20 µL EvoA and re-wetted with 100 µL EvoA. All centrifugation steps were performed at 700 × g for 1 min.

### SPEC workflow

SPEC tips were prepared by placing one plug of SAX or SCX material (3M Empore) in a pipette tip using a blunt-ended syringe needle. For C18 experiments, commercial Evotip Pure (Evosep Biosystems) were used. Tips were activated with 10 µL 100% DMSO (700×g, 1.5 min), then equilibrated with 20 µL buffer (SAX: 20 mM CAPS, 0.01% DDM; SCX: 50% MeOH, 1% FA, 0.01% DDM; C18: 1% FA, 0.01% DDM) at 700×g for 2 min.

Protein samples diluted in equilibration buffer (1:10 for SAX/SCX) were loaded by centrifugation at 200×g for 10 min. Tips were washed once with 20 µL 50 mM TEAB, 0.01% DDM (SAX/SCX) or 1% FA, 0.01% DDM in ACN (C18) at 700×g for 1.5 min. On-tip digestion used 5 µL digestion buffer (0.25 µg/µL trypsin/LysC in 50 mM TEAB, 0.01% DDM), centrifuged briefly at 100×g for 20 sec. Standard digestion time was 60 min at 37°C, though kinetics experiments used 5–90 min.

For two-tip workflows, peptides from SAX/SCX tips were eluted directly into prepared Evotips Pure by adding 20 µL elution buffer (SAX: 0.01% DDM, 1% FA; SCX: 0.01% DDM, 100 mM CAPS). C18 tips were centrifuged to remove digestion buffer and washed once with buffer A.

### Solution digest

In-solution digestion: Sample (5 µL) was transferred to a 96-well plate (twin.tec LoBind, Eppendorf) and mixed with 5 µL digestion buffer (composition as for SPEC workflow). The plate was sealed, briefly centrifuged, and incubated at 37° C for 1 h unless indicated otherwise. Digestion was quenched with 10 µL 1% FA, 0.01% DDM, and samples were loaded onto prepared Evotips Pure.

In-solution digestion with SDB-RPS cleanup: Digestion was performed with identical lysate input and protease amounts as SPEC but in 40 µL total volume at 37°C. After digestion, samples were quenched with 100 µL 1% TFA in 2-propanol and loaded onto SDB-RPS StageTips. StageTips were washed with 20 µL 1% TFA in 2-propanol, then with 20 µL 2% ACN, 0.2% TFA, and eluted with 60 µL 80% ACN, 5% NH_4_OH. Eluates were dried by vacuum centrifugation, resuspended in 10 µL 2% ACN, 0.2% TFA, and loaded onto prepared Evotips Pure.

For on-top-of-tip solution digest, the lysate and digestion buffer (both same as for SPEC workflow) was added above a SAX-SPEC tip with 20 µL of 50 mM TEAB, 0.01% DDM. After digestion 50 µL of 20 mM CAPS pH 10.5, 50% MeOH, 0.01% DDM was added and the tip centrifuged at 200xg for 10 min. Sample was directly eluted onto a prepared EvoTip Pure with 20 µL of 1% FA, 0.01% DDM.

### Protein Aggregation Capture (PAC) digest

PAC digestion was performed similarly as described before [cite, Batth]. Hydroxyl beads (Resyn Biosciences) were washed 2x with ACN. Per sample, 5 µL beads (100 µg) were resuspended in 45 µL ACN. Lysate (5 µL, same as for other digestion types) was added, mixed by pipetting three times, and incubated for 10 min at room temperature for protein aggregation capture. Beads were washed once with 100 µL of ACN. After complete ACN removal, 10 µL digestion buffer (0.125 µg/µL trypsin/LysC in 50 mM TEAB, 0.01% DDM; same total protease amount as other methods) was added and samples were incubated at 37°C for 1 h unless indicated otherwise. After digestions samples were quenched with 20 µL 1% FA, 0.01% DDM and loaded onto prepared Evotips Pure.

### SPEC fractionation

For SPEC based fractionation, SAX SPEC was performed as described above. Instead of a single elution peptides were eluted in five fractions. The first four fractions were based on 20 mM TEAB pH 8.5, 0.01% DDM with varying salt concentrations of 0, 50, 100, and 300 mM NaCl. Elutions were performed with 50 µL directly into a separate prepared EvoTip Pure. The final elution was 20 µL of 1% FA, 0.01% DDM. Fractionations shown were performed with 2.5 µg total lysate.

### Mouse tissue preparation

Animals used were bred for scientific purposes, and the research in this project does not involve experiments on animals (as defined by law). All animals were sacrificed by CO2 euthanasia prior to removal of tissue in accordance with the European Commission Recommendations for the euthanasia of experimental animals (Part 1 and Part 2). Breeding, housing, and euthanasia of the animals are fully compliant with all German (i.e., German Animal Welfare Act) and EU (i.e., Directive 2010/63/EU) applicable laws and regulations concerning care and use of laboratory animals.

Eight-week-old male C57BL/6J mice were euthanized and perfused (cardiac) with rinsing solution (100 mM Tris pH 8.5, 200 mM NaCl, 20 mM EDTA); organs were removed, washed, and fixed overnight in 4% paraformaldehyde before paraffin embedding. Soleus and EDL muscles were freeze-dried overnight; single fibers were pulled, pooled by type, and stored at -65°C.

FFPE samples (∼300 µg) were deparaffinized by sequential incubation in n-heptane (2×) and methanol (2×) at 30°C. Deparaffinized tissue or muscle fibers were resuspended in lysis buffer (50 mM TEAB pH 8.5, 2% SDC, 40 mM CAA and 10 mM TCEP), boiled for 5 min at 95°C, tip-sonicated (10 pulses, 5 sec on/5 sec off, 20% duty cycle), and boiled again. Protein concentration was determined by tryptophan fluorescence.

### Fluorescence imaging of SPEC tips

To track proteins and enzymes during SPEC processing, both were fluorescently labeled. K562 lysate (40 µg) was labeled with 20 µg Alexa Fluor 750 NHS Ester (Thermo Fisher Scientific, 10 mg/mL in anhydrous DMSO) for 1 h at room temperature, giving a final labeling concentration of ∼0.95 µg/µL at an estimated 20:1 molar ratio, then quenched with Tris-HCl pH 8.5. Enzymes (trypsin, LysC) were labeled with Alexa Fluor 647 NHS Ester (Thermo Fisher Scientific, 10 mg/mL in anhydrous DMSO) at a final labeling concentration of ∼0.45 mg/mL at a 20:1 molar ratio. Fluorescence imaging was performed on an Axioscan 7 (Zeiss) with Colibri 7 LED source using a 5× NA 0.8 Plan-Apochromat objective, acquiring 647 nm (enzyme) and 750 nm (protein) channels.

### mTRAQ labeling

For on-tip labeling, peptides were digested on SAX SPEC tips as described, except the equilibration and loading buffer was changed to 50 mM Na_2_CO_3_ pH 10.5, 50% MeOH, 0.01% DDM to eliminate nucleophiles that could quench the mTRAQ reagent. Before elution, digestion buffer was spun through at 700xg for 0.5 min. mTRAQ reagent (SCIEX) was diluted to a final concentration of 0.025 U/µl in 50% ACN, 50 mM TEAB pH 8.5, added directly to the tip, centrifuged briefly at 100×g for 20 sec and incubated for 60 min at room temperature. Labeled peptides were then eluted onto C18 Evotips. Labeling efficiency was calculated by comparing labeled to 18 unlabeled peptide intensities in library-free DIA-NN search.

### Blood collection

Whole blood was collected by venipuncture into EDTA-containing tubes (9 mL). The study was approved by the Ethics Committee of LMU Munich (Reg. No. 17-012). Informed consent was obtained from all human subjects and confirmed that the experiments conformed to the principles set out in the WMA Declaration of Helsinki and the Department of Health and Human Services Belmont Report. The tubes were centrifuged at 2000 × g for 10 min to separate plasma from the cellular components. The plasma fraction was carefully removed, aliquoted, and stored at – 80 C until further use.

Neat plasma was diluted to 10 µg/µl in 50 mM TEAB pH 8.5, 40 mM CAA, 10 mM TCEP and 1% SDC, boiled for 10 min at 98°C, sonicated (10 pulses, 5 s on/5 s off, 20% duty cycle), boiled again and stored at -80°a C.

### Plasma glycoproteomics

Neat plasma (5 µg) was processed with SPEC and solution-based digests. For proteome analysis, 500 ng of digest was loaded directly onto Evotips. For glycopeptide enrichment, the remaining digest was applied to cotton HILIC StageTips (cite, Jager / Selman).

Cotton HILIC StageTips were prepared as previously described. A single plug of glass microfiber (GF/C grade, 1.2 µm pore size, Whatman) was placed in a 200 µL pipette tip using a blunt-ended syringe needle. Cotton pads (Demak’Up) were separated and rice-grain-sized balls were pushed into the prepared tips above the glass fiber plug using a blunt-ended syringe needle.

For glycopeptide enrichment, StageTips were equilibrated by washing with 20 µL 80% ACN, 1% TFA and centrifuging at 200 × g for 1 min. Samples were diluted to 80% ACN with 1% TFA in ACN (final volume ∼100 µL) and loaded onto Cotton HILIC StageTips by centrifugation at 50 × g for 5 min. StageTips were washed twice with 20 µL 80% ACN, 1% TFA (200 × g, 1 min). Glycopeptides were eluted directly into prepared Evotips Pure by two additions of 50 µL 2% ACN, 0.2% TFA with centrifugation at 200 × g for 1 min.

### diGly enrichment

diGly enrichment was performed using the PTMScan HS Ubiquitin/SUMO Remnant Motif Kit (Cell Signaling Technology). RPE-1 lysate (5 µg) was digested either by in-solution or SAX-SPEC as described above. For SAX-SPEC samples, peptides were eluted with 20 µL 1% FA, 0.01% DDM, 625 mM NaCl.

For binding, in-solution digest (10 µL) was diluted with 25 µL binding buffer (20% ACN, 100 mM MOPS pH 7.4, 500 mM NaCl, 0.2% NP-40), 5 µL 1 M TEAB pH 8.5, and 10 µL water (50 µL total). SPEC eluate (20 µL) was diluted with 25 µL binding buffer and 5 µL 1 M TEAB pH 8.5 (50 µL total). Samples were incubated with 2.5 µL anti-diGly magnetic beads for 2 h at 4°C with shaking at 1,200 rpm in a 0.5 mL 96-deep well plate (Eppendorf LoBind).

Beads were magnetically separated and washed four times with 100 µL washing buffer (10% ACN, 50 mM MOPS pH 7.4, 250 mM NaCl, 0.1% NP-40) and twice with 100 µL ice-cold water. Enriched diGly peptides were eluted by two additions of 50 µL 0.15% TFA with 5 min incubation at room temperature and 1,200 rpm, eluting directly onto prepared Evotips Pure.

### LC-MS/MS analysis

Samples were analyzed on an Evosep Eno LC coupled to an Orbitrap Astral Zoom (Thermo Fisher Scientific). The Whisper Zoom 80 gradient was used with an Aurora Rapid column (5 cm × 75 µm, 1.7 µm C18; IonOpticks) at 60 °C unless mentioned otherwise. MS1 spectra were acquired at 240,000 resolution (380–980 m/z) with 500% AGC and 3 ms maximum injection time. Narrow-window DIA used 3 Th windows with 6 ms injection time and 25% HCD collision energy.

SPEC kinetic samples were acquired using the Evosep Eno 500 SPD gradient on the EV1182 column (4 cm x 150 µm, 1.9 µm C18; Evosep) at 40 °C using a method with 100 variable windows and 5 ms max injection time for the MS2 scan instead of the described above method.

Glycopeptides were analyzed using the Whisper Zoom 40 gradient with an Aurora Elite column (15 cm x 75 µm, 1.7 µm C18; IonOpticks) at 60 °C and a top50 DDA method with an MS scan at 240,000 resolution (850-2000 m/z) with 50 ms max injection time and a dynamic exclusion time of 15 s. Precursors from charges 2-8 and a minimum intensity of 5000 were isolated with 1.6 m/z isolation width with a 10 ms max injection time and fragmented at 30 % HCD collision energy before analysis in the Astral analyzer with an MS2 mass range set to 150-2000 m/z.

Di-Gly peptides were analyzed identically to the base method except the DIA windows where 150 variable windows from 380-1380 m/z with a max injection time of 6 ms were used^25^.

## Data analysis

Raw files were searched with DIA-NN v2.2 in library-free mode against the UniProt human canonical database (May 2024) or a combined human, S. cerevisiae, Arabidopsis canonical database (Nov 2025)^26^. Settings: 2 missed cleavages, 10 ppm mass accuracy, 5 ppm MS1 accuracy, scan window 6, gene-level protein inference, single-pass neural network classifier with match between runs enabled. Labeled peptides with mTRAQ were searched as described earlier with the same tolerances.

Enriched glycopeptides were analyzed with Glycan Finder v2.5 (Bioinformatics Solution Inc.) with the following settings: precursor mass error tolerance 10 ppm, fragment ion mass error tolerance 20 mDa, glycan fragment ion mass error tolerance 20 ppm, isotopic shift -1 to 3, specific digest mode, 1 missed cleavage, carbamidomethylation (C) as fixed modification, N-term acetylation and M oxidation as variable modifications, 2 maximum variable PTM per peptide, peptide length 6-45, N-linked database with 1867 entries, max fucose count 2, report filtered to 1 % FDR for glycopeptides. Protein and summary csv files from Glycan Finder and parquet report files from DIA-NN were analyzed in Python 3.11 with custom scripts.

For diGly (KGG) enrichment data, raw files were searched using the ‘directDIA+’ workflow in Spectronaut v20.1 against the human reference proteome. Standard settings for a tryptic sample were chosen. PTM localization mode was activated with cysteine carbamidomethylation as a fixed modification and protein N-terminal acetylation, methionine oxidation, and GlyGly (K) as variable modifications. Tabular data were exported using the ‘BGS Factory Report’ schema with additional columns: ‘EG.PrecursorID’, ‘PEP.PeptidePosition’, ‘EG.PTMAssayProbability’, ‘PG.Genes’, and ‘PG.ProteinGroups’. Data were parsed with a custom Python implementation of the ‘PeptideCollapse’ algorithm^27^ to collapse multiply modified peptides to unique KGG sites.

## Acknowledgements

We thank our colleagues at the Department of Proteomics and Signal Transduction at the Max Planck Institute of Biochemistry and at Evosep Biosystems for their support and valuable discussions and Bioinformatics Solutions Inc. for providing early access to GlycanFinder.

## Conflict of Interest

M.M. is an indirect shareholder in Evosep Biosystems. S.S. and N.B. are employees of Evosep Biosystems. All other authors declare no competing interests.

## Data Availability

The raw mass spectrometry data have been deposited in the public proteomics repository MassIVE with the identifierMSV000100271_reviewer.

## Code Availability

Analysis scripts are available at [https://github.com/DenysOliinyk3007/SPEC-figures].

## Supplemental Information

**Supplementary Figure 1.**
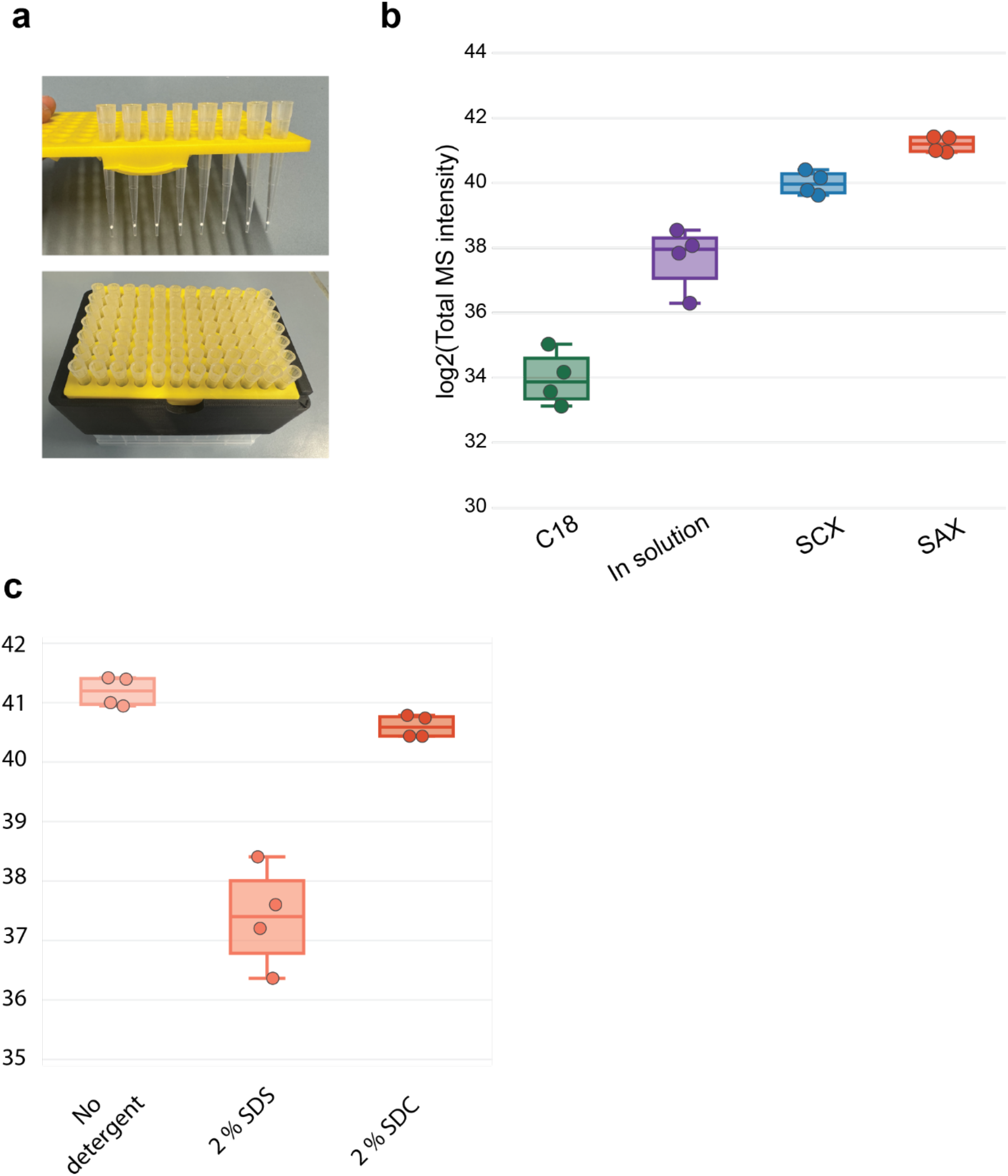
**a** 3D printed adapters for handling SPEC tips. **b** Log_2_ of the total precursor intensity of the same data as **Fig 1c. c** Log_2_ of the total precursor intensity of the same data as **Fig 1d**.

**Supplemental Figure 2.**
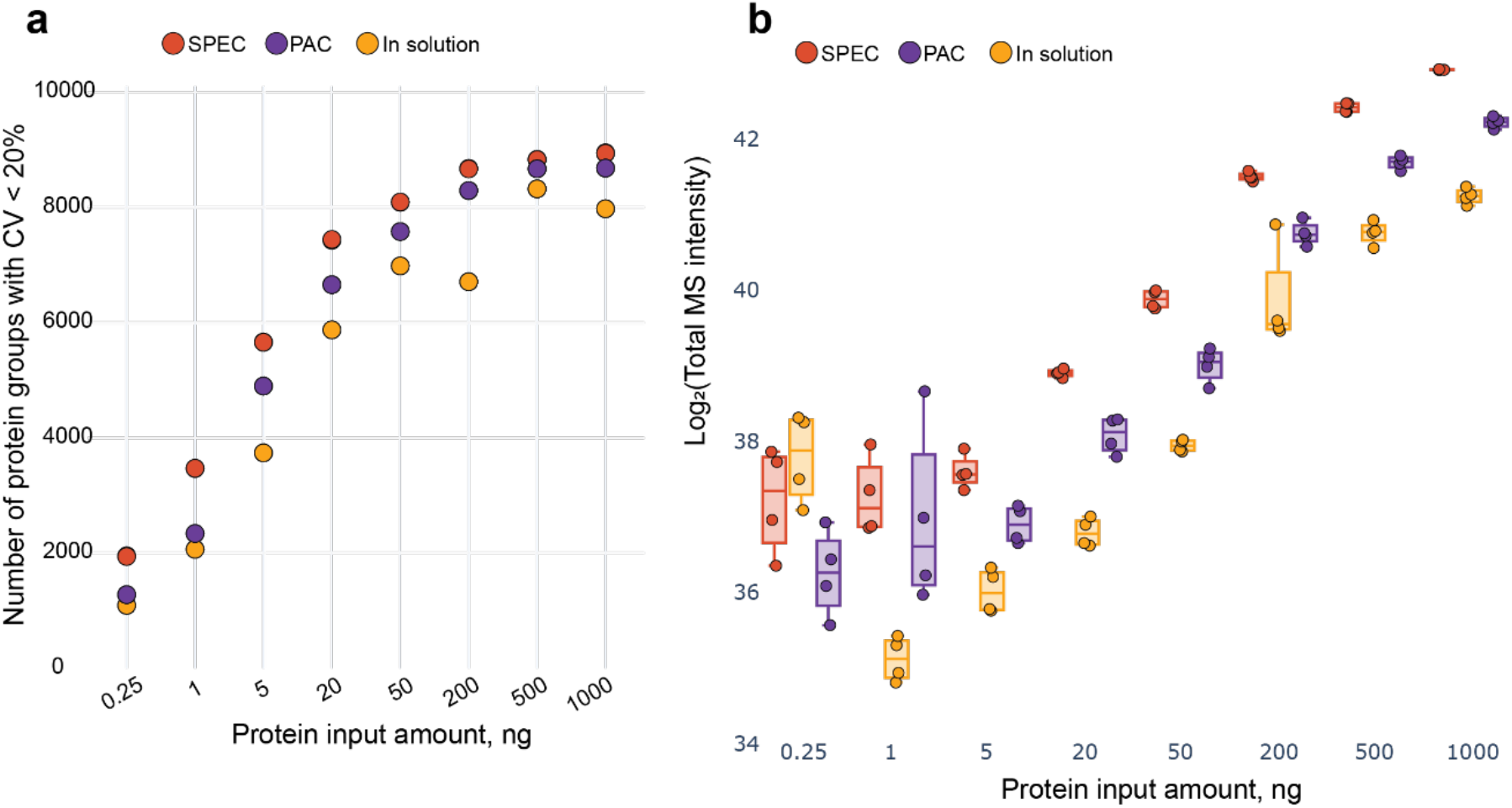
**a** Protein groups with a CV below 20 % from the same data as **Fig 2a. b** Log_2_ of the total precursor intensity of the same data as **a**.

**Supplemental Figure 3.**
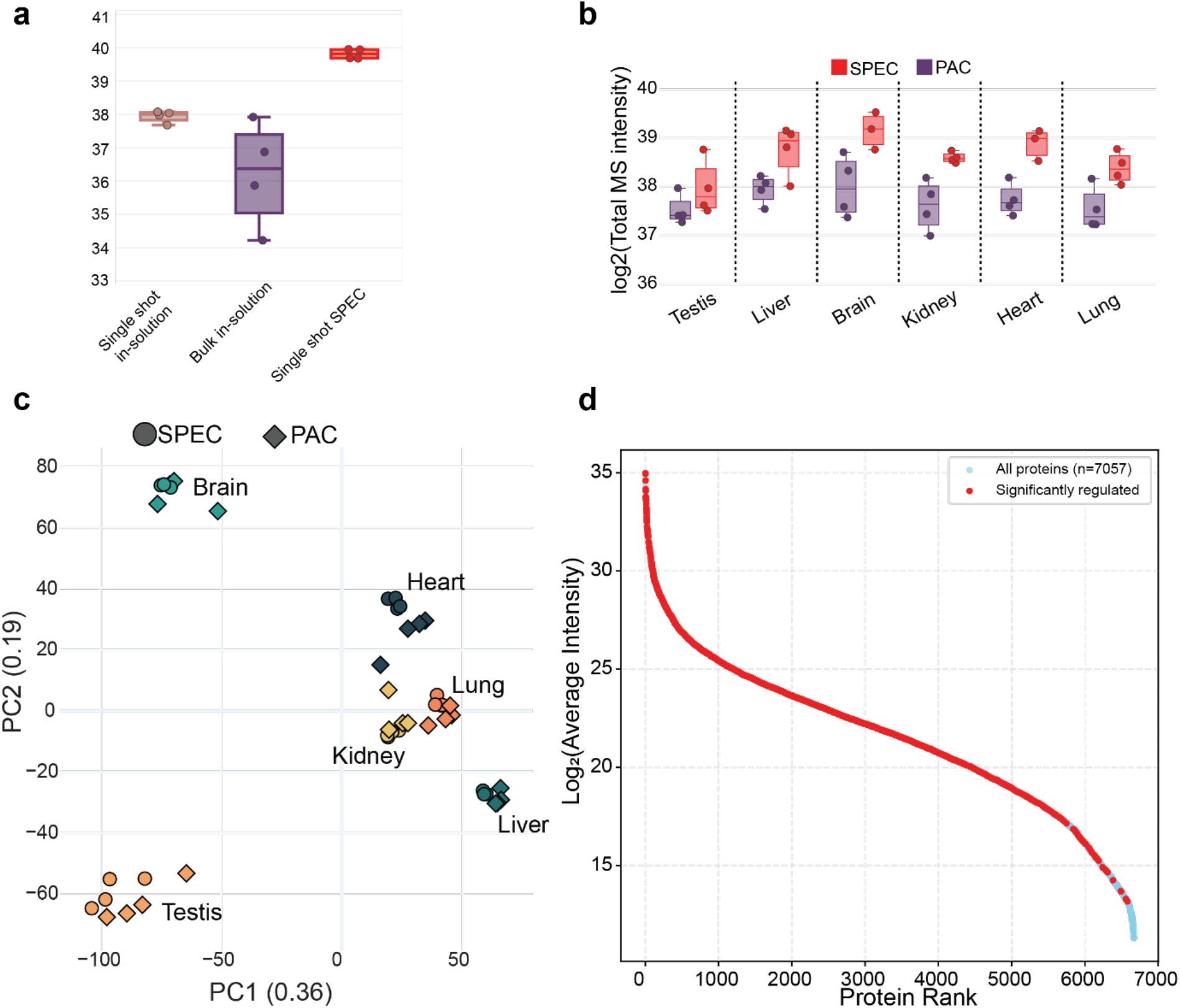
**a** Log_2_ of the total precursor intensity of the same data as **Fig 3b. b** Log_2_ of the total precursor intensity of the same data as **Fig 3f. c** PCA of the data of **Fig 3f. d** Rank plot of the detected proteins of the muscle tissue from **Fig 3d** with significantly regulated proteins highlighted in red.

**Supplementary Figure 4.**
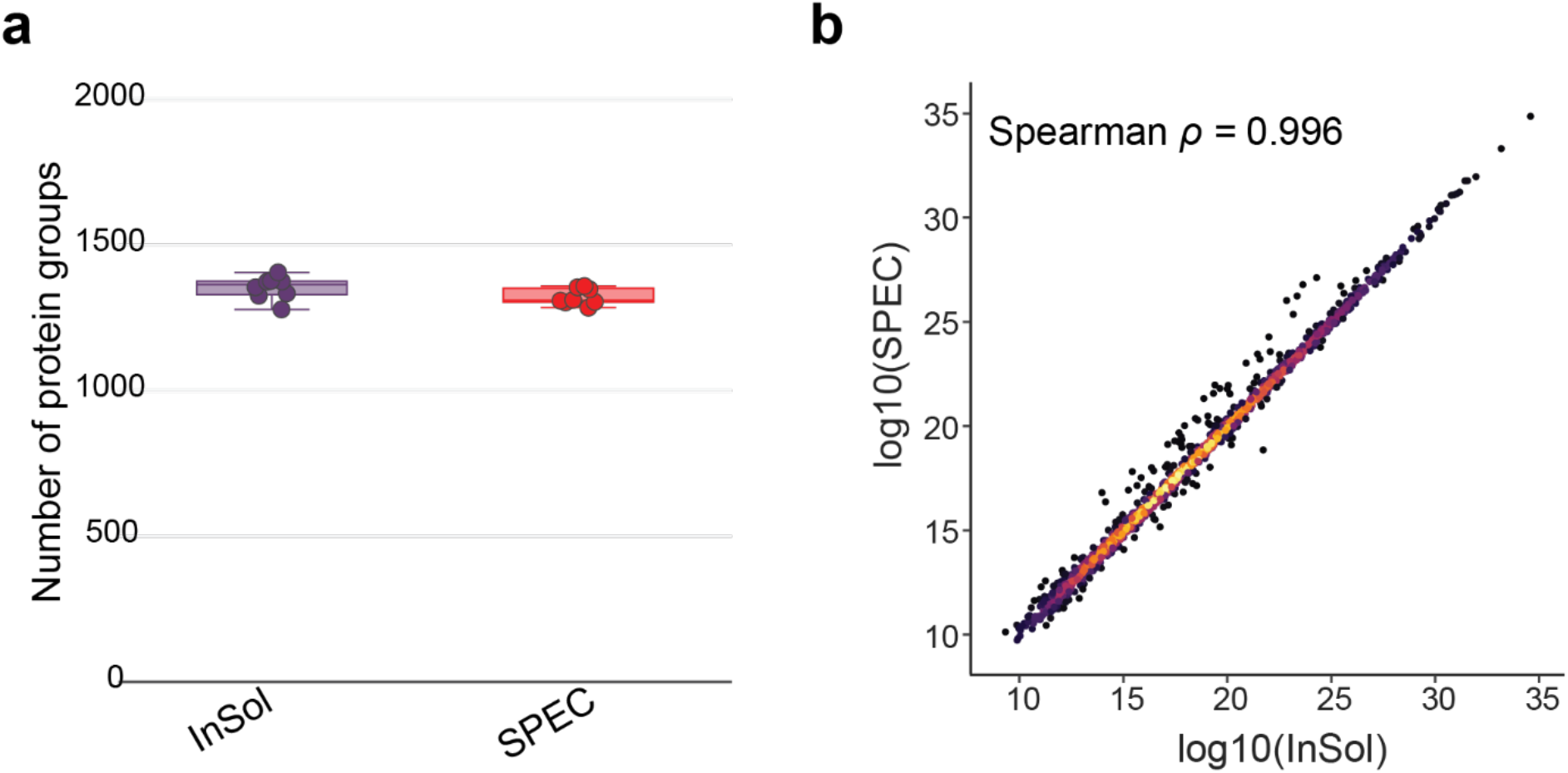
**a** Protein group identifications of the proteome from the same sample prior to enrichment from **Fig 4b. b** Spearman protein rank correlation between the in solution and SPEC plasma digest from **a**.

## Notes

### Competing Interest Statement

Mathias Mann is an indirect shareholder in Evosep Biosystems. S.S. and N.B. are employees of Evosep Biosystems. All other authors declare no competing interests.

### Summary of Updates

This version is supplemented with the newest data

